# The role of specific phosphorylation patterns in the oligomerization of Tau-R4

**DOI:** 10.1101/2024.12.14.628478

**Authors:** Shachar Guy Bressler, Dana Grunhaus, Amit Aviram, Stefan G.D. Rüdiger, Mattan Hurevich, Assaf Friedler

## Abstract

Specific phosphorylation patterns control the activity of multiphosphorylated proteins. In case of the Tau protein, multiphosphorylation leads to the formation of different disease-related condensates and aggregates. Studying the role of these specific patterns at the protein level is crucial for understanding the molecular mechanisms of Tauopathies such as Alzheimer’s Disease. However, due to the extreme difficulty in obtaining recombinant proteins with specific phosphorylation patterns using kinase-based methods, it is practically impossible to study the connection between specific phosphorylation patterns and aggregation events at the protein level. Here we addressed this problem by reducing the system to the peptide level and studying the effect of specific phosphorylation patterns on the condensation and aggregation of a specific domain of Tau, R4 (residues 336-358). To achieve this aim, we have applied advanced methods to synthesize a library of multiphosphorylated peptides derived from R4. We showed that specific phosphorylation patterns stringently control the formation of Tau aggregates and condensates. Phosphorylation of Ser341 promoted aggregation of R4 while phosphorylation of Ser352 promoted its condensation. Interestingly, Ser356 phosphorylation inhibited both processes, which can be overridden by double-phosphorylation at Ser341/Ser352. Differences between the microenvironments of the phosphorylated residues lead to their different effects on R4 aggregation upon phosphorylation. Our results show that working at the domain level using advanced peptide synthesis methods is a highly useful and practical way to provide valuable information about the effects of post translational modifications on protein activity.

## Introduction

Phosphorylation of serine and threonine is one of the most abundant posttranslational modifications (PTM) in eukaryotes^1, 2^. Phosphorylation has a key role in regulating protein-protein interactions (PPI) and self-assembly of monomeric protein to oligomers^3-5^. The most prevalent phosphorylation sites occur on serine and threonine, and most phosphoproteins are phosphorylated at multiple sites^4, 6^. Proteins can oligomerize to different types of assemblies, ranging from coacervates formed via liquid-liquid phase separation (LLPS) to aggregates of different types such as amorphous, native-like, or amyloid fibrils^7^. In cells, coacervates function as membrane-less organelles, which have several possible functions such as bioreactors that catalyze protein interactions and cellular processes^8-10^. Aggregation, however, is frequently associated with pathological pathways that lead to disease^11^.

Different phosphorylation patterns can regulate protein activity and dictate which pathway the protein will undergo. They regulate PPI and may control the oligomerization type of proteins, selecting between aggregation and condensation^3, 12^. The existence of multiple phosphorylation sites in proteins means that there are many possible phosphorylation patterns, each with a unique effect on oligomerization. The ideal way for studying the effect of each such pattern is by obtaining a library of multiphosphorylated derivatives of the protein of interest, each with a unique and specific phosphorylation pattern. Such a library would cover all relevant patterns. However, obtaining such a diverse multiphosphorylated protein library in sufficient quantities for extensive biophysical, biochemical and structural studies is not feasible.

*In-vitro* phosphorylation of recombinant proteins by kinases lacks the specificity to provide a library of individual proteins, each with a different pattern. Many kinases are not totally selective, and for some phosphorylation sites it is not even known which kinase phosphorylates them^13^. Semi-Synthetic and synthetic approaches based on native chemical ligation (NCL) were developed to access proteins as a way to overcome the problems associated with expression and enzyme selectivity^14-20^. However, obtaining proteins with defined phosphorylation patterns via synthetic approaches still sets a huge challenge. The synthesis of each protein involves a huge amount of work and time for the production of a small quantity that is, arguably, insufficient for performing the required studies. This limits the use of synthetic proteins as practical tools to study phosphorylation patterns.

An alternative way to assess the effects of such patterns is by using glutamate residues for replacing the phosphorylated residues. These point mutations can be done *in vitro* or *in cells* ^21, 22^. However, those mutated proteins do not offer a genuine model for the native ones. There is a huge difference between a phosphate group and a carboxylic group. Replacing a phosphoserine or phosphothreonine by glutamate retains only the negative charge at the specific site but overlooks all the unique chemical and physical properties of the phosphate group^21^. A phosphate has two negative charges at physiological pH, while glutamate has only one. Moreover, the pKa of phosphate is 5.58, while the pKa of glutamate is 4.25^23^. The phosphate also occupies much larger volume that is derived from its tetrahedral geometry compared to the planar carboxylate group. These factors govern the distinctive interplay between the phosphorylated residue and its surroundings, dictating the role of the phosphate at each specific microenvironment within the protein. Comparative studies using negatively charged residues such as Glu showed that Glu substitution of the phosphorylated Ser does not necessarily result in the same protein structure or function. ^24 21^ This was demonstrated for example for α-synuclein, where phosphorylation of Ser219 inhibited its fibrillation, while the mutation of S219 to D or E did not. ^24 21,25^. Using synthetic phosphorylated peptides offer an alternative to study the effect of patterns on interactions and oligomerizations, which overcomes the above problems. Peptides have several advantages in bridging the gap between the authentic yet inaccessible proteins to the much-needed biophysical and biochemical input. First, synthesis of peptides can provide each specific pattern at high purity, specificity and reliability. Second, synthetic processes guarantee that the peptides can be obtained reproducibly and at a sufficient quantity. Third, peptides can be labeled by different chemical tags to enable analysis via diverse biochemical assays.

Synthesizing multiphosphorylated peptides is difficult since protected phosphorylated Ser\Thr used in the synthesis tend to undergo elimination under the standard Fmoc deprotection condition. These protected phosphorylated building blocks are also very bulky, making the couplings less efficient. Hence, obtaining a library of long multiphosphorylated peptides derived from a specific protein domain, in sufficient quantities and purities is practically impossible. It is time consuming and even if some peptide is obtained, its synthesis is extremely inefficient. This makes accessibility to phosphorylated peptides a limiting factor in studying phosphorylation-controlled biological processes such as protein aggregation. To overcome these limitations, we have previously developed a new method for the rapid synthesis of multiphosphorylated peptides in large amounts and high purities^20, 26-28^. The method is based on a combination of high temperature and fast mixing for accelerating each coupling step while minimizing the formation of side products and maximizing the efficiency^26^. Using this method, we demonstrated the synthesis of peptides containing up to eight phosphorylation sites. The above strategy is ideal for synthesizing libraries of multiphosphorylated peptides. These libraries can then be used for systematic studies of the effect of specific phosphorylation patterns on the biological activity of proteins. Here we applied our method for studying the aggregation processes of the Tau protein^29, 30^.

The Tau protein forms amyloid fibrils in neurons as part of the pathology of neurodegenerative diseases (ND). *in vivo*, fibrils of Tau are hyperphosphorylated^31^, an event that is linked to aggregation and condensate formation by the protein^13, 32-35^. However, the role of the specific phosphorylation patterns in controlling the initial molecular events leading to amyloid aggregates or condensates is mostly unclear. Tau is a highly disordered protein that is composed of an N-terminal domain, a proline-rich domain, microtubule binding domain (RD) with five homologous repeats (R1-R4, R’) and a C-terminal part (Figure 1A). In Tauopathies such as Alzheimer’s disease (AD) and chronic traumatic encephalopathy (CTE), Tau undergoes aggregation into amyloid fibrils via its RD. Fibrils from AD and CTE patients share a hairpin with a hydrophobic core that is formed by the fourth repeat (R4) of Tau-RD (Figure 1B). In fibrils, R4 has three beta-sheets, interconnected by disordered linkers. in addition, Tau-RD is essential for the condensation of Tau^35, 36^. Although it is known that phosphorylation of the full-length Tau initiates aggregation and condensation^13, 37^, there is almost no information about the effect of specific phosphorylations, mostly due to the difficulty in obtaining Tau derivatives with specific phosphorylation patterns. For example, single phosphorylations at Ser293 in R2 or Ser305 in R3 promote amyloid fibril formation *in vitro* ^17^. However, the effect of specific phosphorylation patterns on oligomerization of R4 is yet unknown. R4 has three sites that were found to be phosphorylated in ND: Ser341, Ser352 and S356. Each of these serine residues is located at a different chemical environment. Ser341 is located between two charged residues, Lys340 and Glu342, within the disordered linker. Ser352 is positioned next to Gln351 and Lys353 and is located within a loop structure, while Ser 356 is located between two aliphatic residues, Gly355 and Leu357 at the end of the loop of R4. The side chains of Ser341 and Ser352 are facing toward the core of the hairpin, while Ser356 is facing out (Figure 1C).

**Figure 1.**
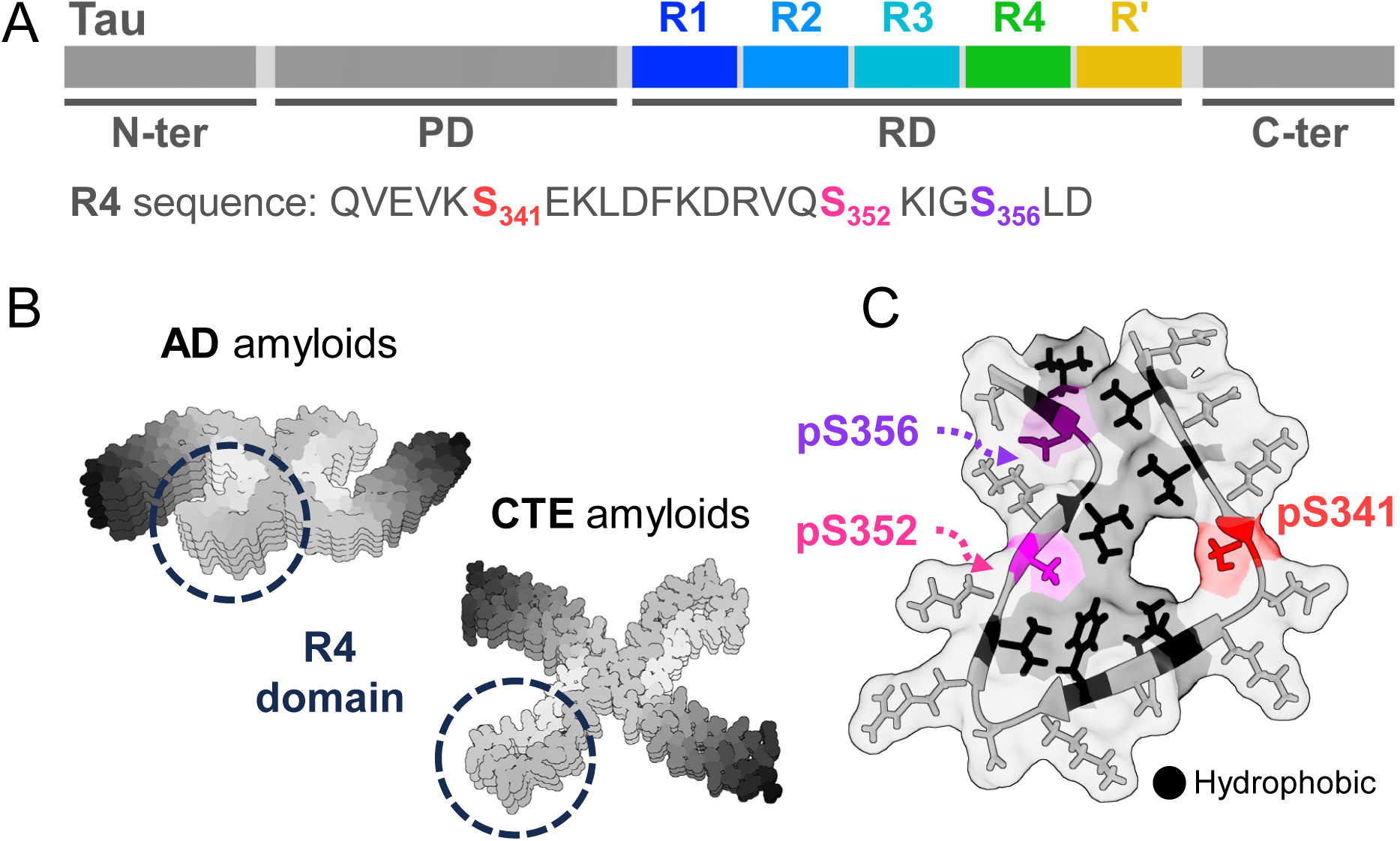
The Tau R4 domain (residues 336-358). Tau 336-358 is derived from the R4 repeat and forms a hairpin structure within native Tau fibrils. (A) Domain structure of Tau: The N-terminal part (N-ter), the proline-rich domain (PD), The repeat domain (RD) with its five repeats R1-R4, R’ (highlighted), and the C-terminus part (C-ter). The sequence of Tau 336-358 (Serine 341 in red, 352 in magenta and 356 in purple). (B) Cryo-EM structures of Tau filaments isolated from patients of different Tauopathies: AD amyloid filament (PDB ID: 5O3T)^38^, CTE amyloid filaments (PDB: 6NWQ)^39^; Tau 336-358 that adopts a hairpin structure is framed. (C) The hairpin structure that Tau 336-358 forms in AD PHF patients. Serine 341 labeled in red, 352 in magenta and 356 in purple. hydrophobic residues in black.

Herein we harnessed our novel approaches for multiphosphorylated peptide synthesis^12, 19, 20^ to study systematically how specific phosphorylation pattern affect aggregation and condensation of the Tau R4 domain. We designed and synthesized a library of peptides representing the R4 domain and containing all the possible phosphorylation patterns. Then, we determined whether each peptide undergoes condensation or aggregation by fluorescence and electron microscopy. We found that the fully phosphorylated peptide can condensate or aggregate depending on the conditions, while the non-phosphorylated peptide cannot oligomerize at all. Phosphorylation of Ser341 promoted aggregation, while phosphorylation of Ser352 promoted condensation. Our research emphasizes the importance of a methodological study that explores the effects of phosphorylation patterns on oligomerization using phosphorylated peptide libraries. This study utilizes our novel methods for synthesizing phosphorylated peptides, enabling the discovery of new phosphorylation sites that are crucial to initial aggrgation and their intricate interconnections, which may not be identified by other current state-of-the-art approaches.

## Results

### Synthesis of a phosphorylated peptide library derived from Tau 336-358

To study the effect of phosphorylation on the condensation and aggregation of R4, we synthesized a library of peptides with patterns including all combinations of phosphorylations in Ser341, Ser352, and Ser356. The peptides were synthesized using our newly developed Accelerated MPP synthesis^20^, which enabled us to synthesize the library in a short time and in sufficient amounts (See Table 1 for peptide sequences and Figures 1S, 2S for the analytical data of the peptides). The synthesis was performed with a constant overhead stirring of 1200 rpm, at 90 °C. The assembly of each peptide was completed in less than 35 min. Sixteen fluorescent and non-fluorescent peptides were obtained in over 95% purity and multi-mg scale in less than two weeks calculated work. The peptides were then studied for their oligomerization using biophysical analyses as described below.

**Table 1.**
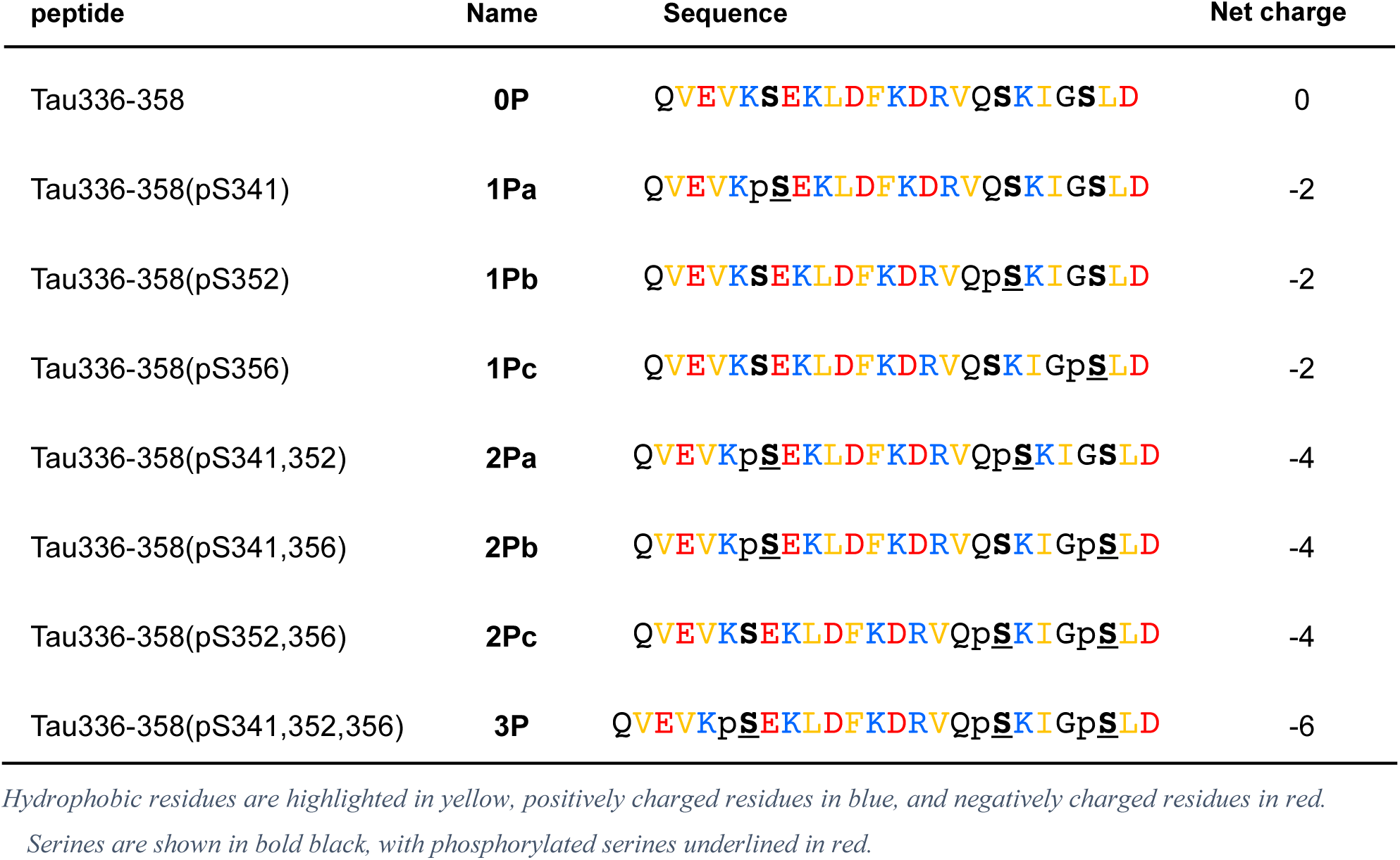
Library of Tau 336-358 and its derived phosphorylated peptides.

| peptide | Name | Sequence | Net charge |
| --- | --- | --- | --- |
| Tau336-358 | **0P** | Q**VEVK**<u>**SEKLDFKDRV**</u>Q**SKI****GS****LD** | 0 |
| Tau336-358(pS341) | **1Pa** | Q**VEVK**p<u>**SEKLDFKDRV**</u>Q**SKI****GS****LD** | -2 |
| Tau336-358(pS352) | **1Pb** | Q**VEVK****SEKLDFKDRV**Qp<u>**SKI**</u>**GS****LD** | -2 |
| Tau336-358(pS356) | **1Pc** | Q**VEVK****SEKLDFKDRV**Q**SKI**Gp<u>**S**</u>**LD** | -2 |
| Tau336-358(pS341,352) | **2Pa** | Q**VEVK**p<u>**SEKLDFKDRV**</u>Qp<u>**SKI**</u>**GS****LD** | -4 |
| Tau336-358(pS341,356) | **2Pb** | Q**VEVK**p<u>**SEKLDFKDRV**</u>Q**SKI**Gp<u>**S**</u>**LD** | -4 |
| Tau336-358(pS352,356) | **2Pc** | Q**VEVK****SEKLDFKDRV**Qp<u>**SKI**</u>Gp<u>**S**</u>**LD** | -4 |
| Tau336-358(pS341,352,356) | **3P** | Q**VEVK**p<u>**SEKLDFKDRV**</u>Qp<u>**SKI**</u>Gp<u>**S**</u>**LD** | -6 |

*Hydrophobic residues are highlighted in yellow, positively charged residues in blue, and negatively charged residues in red.*
*Serines are shown in bold black, with phosphorylated serines underlined in red.*

### Full phosphorylation of Tau 336-358 promotes its aggregation and condensation

To test if the phosphorylation of Tau 336-358 affects its aggregation and condensation, we first compared the fully phosphorylated (**3P**) and the non-phosphorylated (**0P**) peptides. We performed turbidity assays as a quick screening method to test first if the peptides undergo any type of high-order oligomerization. This may indicate possible condensation or aggregation of the peptides. Mixtures of 5% Fluorescein labeled (FL) **0P** and **3P** with 95% non-labeled peptides were tested at NaCl concentrations of 0, 150, 500 and 1000 mM, and peptide concentrations of 50, 300 and 1000 µM. Testing the dependence of the peptide oligomerization on the ionic strength was performed to examine whether the process is charge-dependent. This is important since the phosphorylation adds extra negative charges to the peptides. **3P** showed high turbidity in 1 mM peptide at all NaCl concentrations. It also showed high turbidity values in 1 M NaCl at all peptide concentrations. In contrast, **0P** showed low turbidity at all tested conditions except 1 M NaCl (Figure 2A).

**Figure 2.**
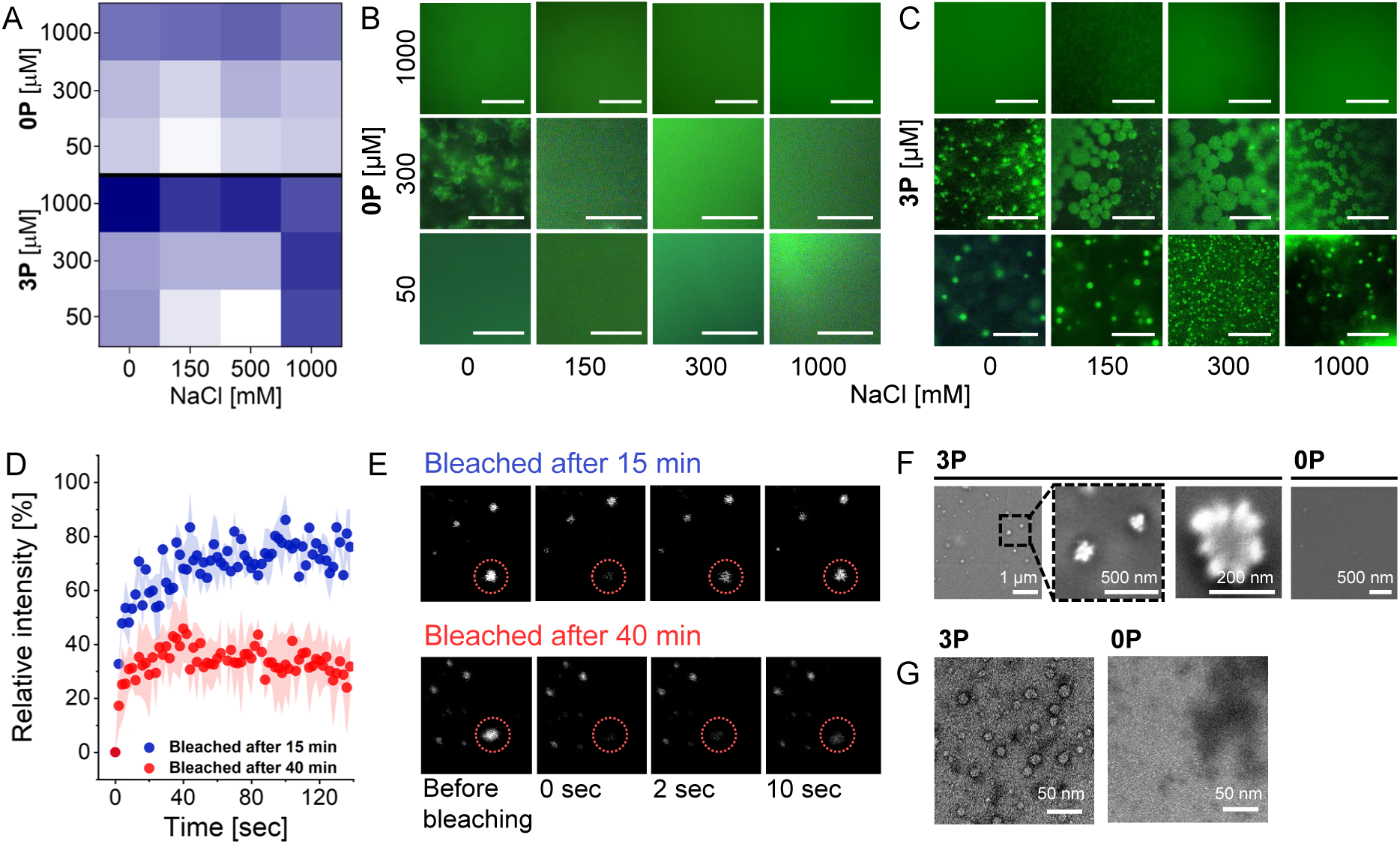
(A) Turbidity assay for initial evaluation of possible oligomerization of **0P** and **3P**. Turbidity was measured in 0, 150, 300 and 1000 mM NaCl concentrations. Darker colors indicate higher optical density that correlates with high order oligomerization (B,C) Fluorescence microscopy images of **3P** and **0P** condensates formed in 0, 150, 300 and 1000 mM NaCl (Y axis) and 50 and 300 µM peptide (X axis) (D) FRAP of **3P** mixed with 5% of FL-**3P** after incubation of 15 min (blue) and 40 min (red). Shown is the time dependence of the relative fluorescence intensity (E) Recovered **3P** bleached condensates (circled), bleached after 15 min incubation on the left and after 40 min incubation on the right, before and during the recovery process. (F) Characterization of **3P** aggregates using SEM microscopy. From left to right: Image of **3P**, followed by zooming in on the flower-like structure and an enlarged image of flower-like aggregates. **0P** did not aggregate. (G) Characterization of **3P** aggregates using TEM microscopy. **3P** showed spherical aggregates. **0P** did not aggregate.

To test if the high turbidity of **3P** resulted from aggregation or condensation, we analyzed the oligomers using fluorescence microscopy. For **3P**, we observed condensates formation in 50 µM and 300 µM of peptide at all NaCl concentrations and aggregation of the peptide at 1000 µM of peptide in all NaCl concentrations (Figure 2B). For **0P** we did not observe aggregation nor condensation of peptide except at very high salt (Figure 2C). This also confirmed that the fluorescein labeling itself did not promote condensation or aggregation.

An important feature of self-assembly processes is their dynamics, meaning the ability of the peptides to diffuse in and out of the condensates. This defines the rate of condensate maturation which is the conversion of condensates into a more viscous gel-like phase ^40^. To evaluate the dynamics of the condensate formation by FL-**3P** and test whether they have similar properties to the condensates of the recombinant Tau^40^, we used fluorescence recovery after photobleaching (FRAP). This method allows monitoring the diffusion of molecules such as peptides in and out of the condensates. Higher recovery means a higher diffusion rate and less viscous condensates. We bleached the entire **3P** condensate areas 15 and 40 minutes after they were formed. ∼80 % of the condensates that were bleached after 15 min recovered after 30 seconds. More mature condensates that were bleached 40 min after formation showed only partial recovery of ∼35 %. In both cases, a *plateau* was reached at approximately 40 seconds (Figure 2D). These results indicate that after 15 minutes the condensates are liquid-like, which allows fast diffusion of **3P** in and out of the condensates. The partial recovery of the mature condensates can indicate a solidification process that led to a more gel-like character (Figure 2D-E).

TEM and SEM images of **3P** taken after 10 days (Figure 2F, 2G) showed that it formed spherical aggregates of 10-60 nm. These units together formed higher-order oligomeric structures with a flower-like morphology. Under the same conditions, **0P** did not aggregate even after this long incubation period.

### The contribution of specific phosphorylations and their combinations to Tau 336-358 aggregation and condensation

To understand the role of each phosphorylation site in controlling the condensation and aggregation of Tau 336-358, we characterized the ability of the mono and di-phosphorylated peptides to form condensates at NaCl concentrations of 0, 150, 300 and 1000 mM (Figure 3A, B) using fluorescence microscopy. Mixtures of 5 % fluorescein-labeled and 95 % of non-labeled peptides were prepared. **1Pa**, **1Pb** and **2Pa** formed condensates, while **1Pc**, **2Pb** and **2Pc** did not form condensates. Like **3P**, **1Pb** and **2Pa** formed condensates in 0, 150, 300 and 1000 mM NaCl. In contrast, **1Pa** formed condensates only at lower salt concentrations of 0 and 150 mM. **2Pb** is the only peptide that underwent condensation only at 1000 mM NaCl. These findings suggest that: 1) Phosphorylation is required for condensation of Tau 336-358; 2) Specifically, phosphorylations of Ser341 and Ser352 facilitate condensation; 3) Phosphorylation of Ser356 inhibits condensation. A closer investigation of the size of the condensates showed that while **1Pa** had an average diameter of ∼3.5 µm, **1Pb** formed condensates with diameters that ranged between 4 to 12 µm. The largest condensates of **1Pb** were in the same range as the mean diameter of **2Pa** ∼12 µm, and **3P** ∼12.3. condensates of **1Pa**, **2Pa** and **3P** were uniform in their size, with relatively small standard deviations, while **1Pb** showed a large range of condensate size (Figure 3C). The intensity of the fluorescence in each condensate was examined since it can indicate peptide concentration. We showed that while the peptide concentration of **1Pb**, **2Pa** and **3P** was homogeneous across the condensate droplets. The edge of **1Pa** condensates had higher fluorescence intensity (Figure 3D) suggesting that peptides accumulated on the surface of the condensate, which can stop the growth of the condensates to give smaller entities.

**Figure 3.**
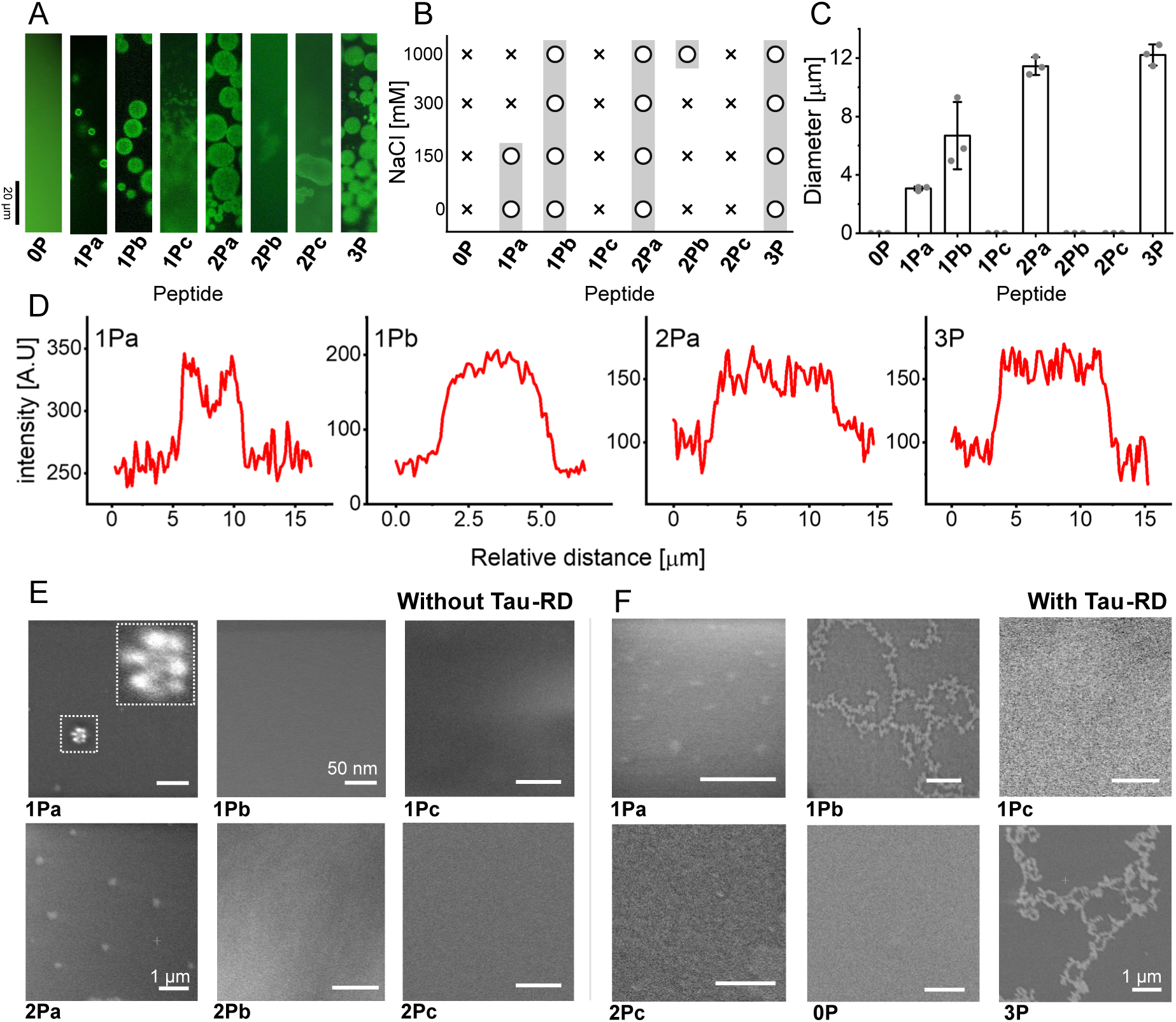
(A) Fluorescence microscopy of the peptide library. Only **1Pa**, **1Pb**, **2Pa** and **3P** made condensates. (B) The dependence of peptides condensates on ionic strength. ‘O’ with grey background: condensates formation; ‘X’: No condensates formation. X axis is the peptide library, Y axis is NaCl concentration in mM. (C) The maximum diameter size of the condensates of the different peptides. (D) Cross-section analysis of condensates of the peptides that underwent condensation. X axis is the position within the cross-section of the condensate in µm. The 0 µm position was defined as the starting point of the measurement. Y axis is the intensity of each position. **1Pa** has higher fluorescence intensity around the condensate edges while in the other peptides, the intensity is unform. (E) SEM of mono- and di-phosphorylated Tau336-358 peptides without Tau-RD (F) SEM of **0P**, **1Pa**, **1Pb**, **2Pc** and **3P** Tau336-358 peptides with Tau-RD. Scale bar = 500 nm.

SEM was used to characterize the aggregates formed by the phosphorylated peptides after 10 days of incubation (Figure 3E). **2Pa**, **1Pb**, **1Pc**, **2Pb** and **2Pc** did not aggregate. Like **3P**, **1Pa** formed spherical aggregates with flower-like morphology. These results suggest that phosphorylation of Ser341 contributes to the aggregation behavior of Tau 336-358, while phosphorylation of Ser352 and Ser356 did not contribute to the phenomena.

### The effect of specific phosphorylations and their combinations on Tau-RD aggregation

Since it is technically impossible to study the effect of phosphorylation patterns at the protein level via a library of phosphoproteins. We decided to study the effect of phosphorylation patterns on Tau oligomerization by testing the effect of the phosphorylated peptide library on aggregation the Tau-RD domain (244-372), which is responsible for its amyloid aggregation. *In vitro* Tau forms amyloids via its MTBR only at the presence of an inducer such as heparin, which is highly negatively charged. Here, we tested if the phosphorylated peptides have a similar effect, due to their negative charges. When Tau-RD was incubated for 10 days with **3P**, spherical aggregates with a diameter of 20-60 nm were formed, interconnected by larger amorphous aggregates. Incubation of the Tau-RD alone or with **0P** for 10 days did not result in aggregation (Figure 3F). Suggesting that aggregation is indeed dependent on the interactions between **3P** and Tau-RD.

To test the effects of the specific phosphorylation patterns at the protein level, **0P**, **1Pa**, **1Pb**, **1Pc** and **2Pc** were incubated with the recombinant Tau-RD for 10 days at 37 °C and the samples were tested using SEM (Figure 3E). **3P** and **1Pb** induced Tau-RD aggregation, resulting in aggregates with string-like morphology. **1Pa**, **1Pc**, and **2Pc** incubation with Tau-RD did not result in oligomer formation.

### The three phosphorylation sites differ in their chemical environments

Charge and steric hindrance alone cannot explain the different effects of the three phosphorylations on the oligomerization of Tau R4. We used the PEP-FOLD3 predictor to examine the conformations and secondary structures of Tau 336-358 ^41^. The most probable 29 models that were calculated and generated (Figure 2S) shared the same closed hairpin-like conformation as was found in AD amyloids (Figure 5A,B). All these models contain a combination of disordered and structured domains (Figure 1C). A notable difference is that according to the models, the Tau 336-358 fragment has a helical domain whereas the same region in the full-length protein has a beta-sheet structure (figure 5A, B). Indeed, peptides usually adopt helical rather than a β-sheet structures since the stabilizing partner strands are missing. In all the calculated models, Ser341 is in the disordered region of the peptide, from Q356 to Phe346. Ser352 is in the helical domain that includes 7 residues from Lys347 to Lys353, and Ser356 is located in the IDR tail from Ile354 to Asp358 (Figure 4A), similar to their locations in the AD amyloid structure. The CD spectrum of **0P**, had a minimum at ∼200 nm and ∼225 nm, corresponding to disordered and helical conformations (Figure 4B). 0P and 3P had similar CD spectra suggesting they obtain the same secondary structure (Figure 4B). Using the PEPstrMOD server ^42^, we generated a *de novo* structure prediction of Tau 336-358 phosphorylated at all three p-sites. The results show that the phosphorylations do not affect the secondary structure (Figure 4C). Thus, we added the phosphate groups to the backbone of 0P using a PyTM plugin (version 1.8.4.0)^43^. Using PyMol measurement wizard^44^, we found that Ser341 and Ser356 are more solvent-exposed than Ser352 (Figure 4D). Calculated distances between the p-site and their nearest entity showed that: 1) pSer352 may interact with the amine group on the backbone of Val337 via hydrogen bond of 1.5 Å, 2) pSer341 had a 4.3 Å distance from the most adjacent positive charged Lys351, and 3) pSer356 had a 4.7 Å distance from the adjacent amide of the side chain of Gln351 (Figure 4E). Altogether, pSer 352 is the least exposed of all three. pSer341 and pSer356 are more exposed, and in contrast to pSer352, do not have significant interactions with their surroundings.

**Figure 4:**
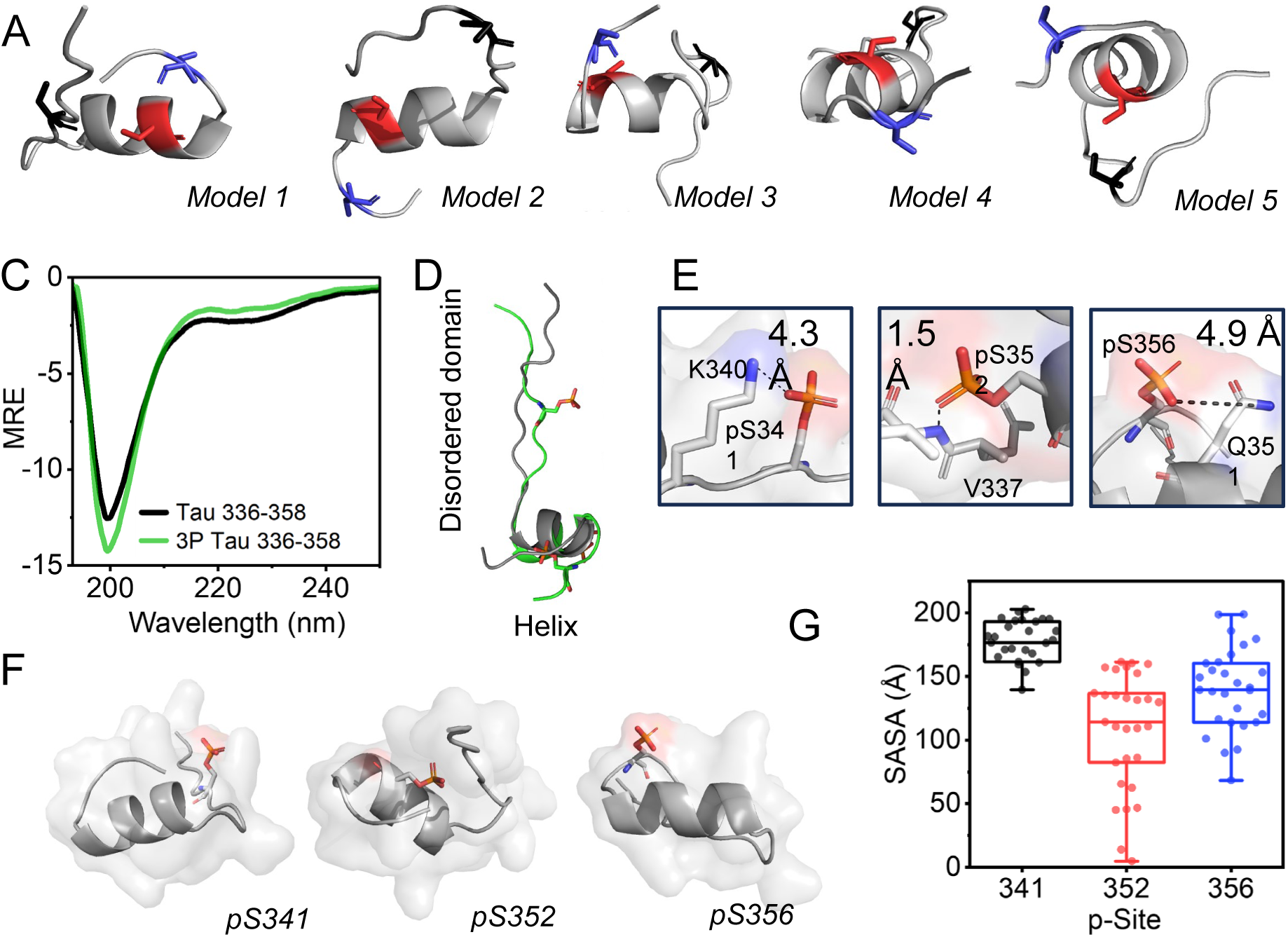
(A) The five most probable models calculated using PEP-FOLD3 ^41^ show that Tau 336-358 has an IDR tail, a helical domain, and a disordered region. In all models Ser341 (black) is located in the disordered region, Ser352 (red) in the helical domain and Ser356 (blue) in the IDR tail. (B) CD spectra of **0P** (yellow) and **3P** (black) indicate the same secondary structures, comprising of helical and disordered domains. (C) models of **0P** (black) and **3P** (green) as calculated by PEP-FOLD3 and PEPstrMOD ^42^. Phosphate groups are shown in orange. (D) Zoom in on the chemical environment and interactions of each phosphorylation site (Ser341 and 356 shown in model 1, Ser352 in model 2). (E) the position of each phosphorylation site in the Tau peptide, as shown by Pymol for model 1. (F) SASA of each phosphorylation site. boxplots represent the median value and interquartile (25-75% percentiles) of SASA. Each dot represents one of the 29 models that were examined.

**Figure 5:**
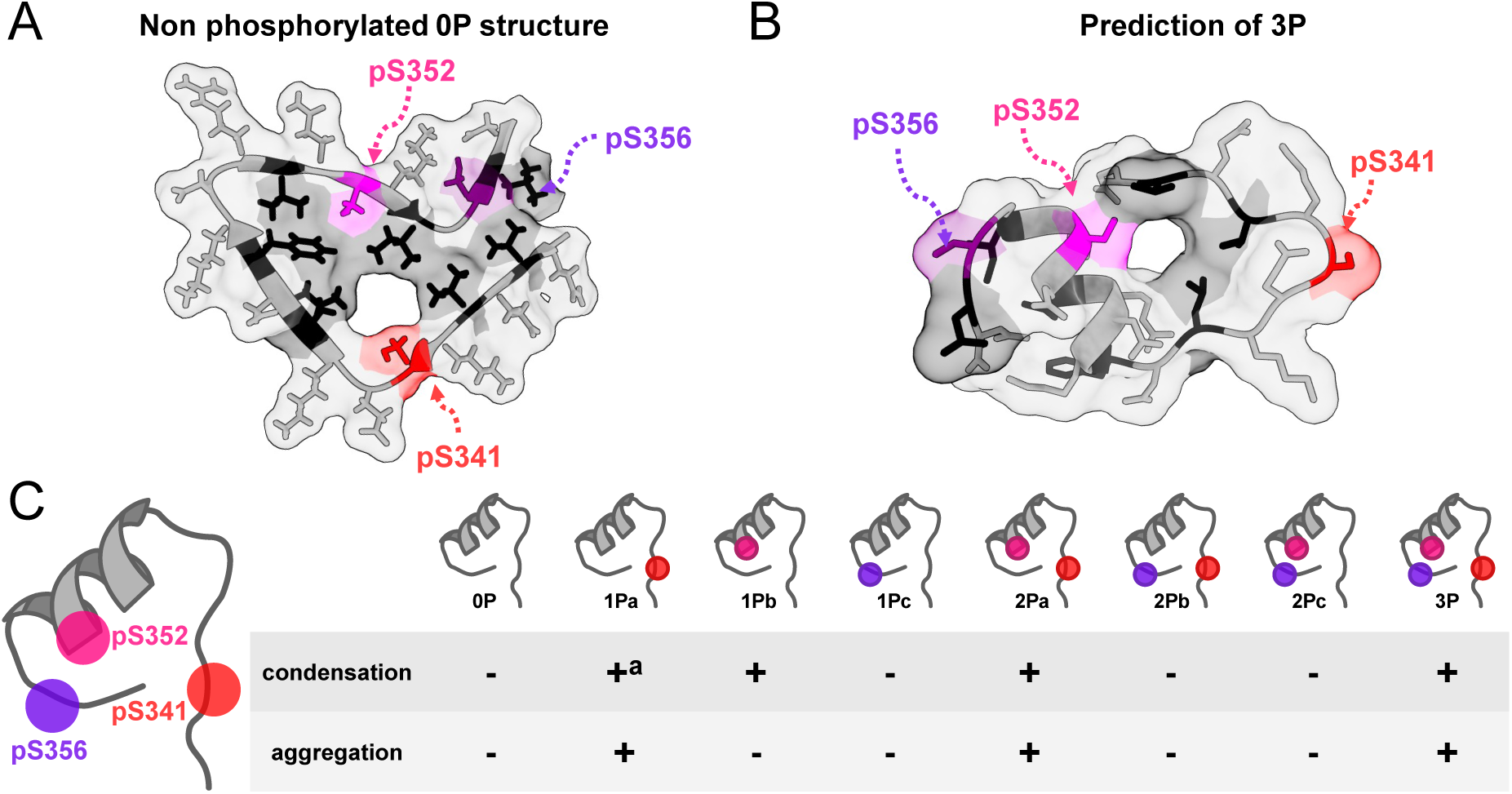
(A) Structure of Tau336-358 **(0P)** as found in amyloids isolated from AD patients (11). (B) Predicted secondary structure and conformation of tri-phosphorylated Tau336-358 **(3P)** as calculated by PEP-FOLD3 (37). Color code for A and B: Hydrophobic groups are black. Serine 341 is labeled in red, 352 in magenta and 356 in purple. (C) Summary of the effects of the different phosphorylation patterns on Tau oligomerization. For condensation, (+) represents condensation in all tested conditions; (+^a^) represents condensation that occurred at certain ionic strength conditions but not all conditions tested; (-) represents peptides that did not undergo condensation. For aggregation, (+) represents the formation of aggregates; (-) represents peptides that did not undergo aggregation.

Using the Pymol software we calculated the solvent-accessible surface area (SASA)^45^ for pSer341, pSer352 and pSer356 in the most probable 29 models of the peptide. The phosphate groups were added by PyTM^43^. to the models that were generated by the predictor. The average SASA of pSer352 was 108±45 Å^2^ while pSer341 had an average SASA of 169±36 Å^2^. pSer356 had an average SASA of 137±36 Å^2^.

## Discussion

In the current study, we combined phosphorylated peptide libraries with an ensemble of biophysical methods to elucidate the specific effects of phosphorylation patterns on the different association processes of Tau R4. We found that phosphorylations at specific sites dictate whether the peptides can assemble to form oligomers at all, and which type of oligomerization process they undergo or induce (condensation or amyloid aggregation). Our results highlight the importance of Ser352 as a major phosphorylation site for inducing Tau oligomerization. This result could be obtained only with the help of our synthetic methods enabling the synthesis of phosphorylated peptide libraries.

### The role of the phosphorylation pattern in controlling the oligomerization of the Tau R4 domain

Our results show that the Tau 336-358 peptide (**0P**), representing the non-phosphorylated R4 domain, does not form any type of high-order oligomers by itself and does not induce oligomerization of Tau-RD. Phosphorylation of one or more serine residues within peptides derived from this R4 domain results in significant changes of its oligomerization. The tri-phosphorylated peptide **3P** formed condensates or aggregates, depending on the time and ionic strength, and induced aggregation of the Tau-RD. This indicates that phosphorylation controls both the condensation and aggregation of this domain. These phenomena are unique, as recombinant non-phosphorylated Tau monomers do not tend to aggregate even within days^46^.

The contributions of each phosphorylation within these patterns are not necessarily additive. Phosphorylation of Ser352 activated condensation of R4 as shown by the ability of the **1Pb** peptide to undergo condensation, while phosphorylation of Ser341 activated aggregation of R4 as shown by the aggregation of the **1Pa**. peptide. Phosphorylation of Ser356 prevented any oligomerization of R4 as reflected by the inability of **1Pc** to form condensates or aggregates. The effects of combinations of two phosphorylations were synergistic. **2Pa,** which contained phosphorylations at residues Ser341 and Ser352, generated condensates with similar size and propensity to **1Pb** containing a single phosphorylation at Ser352. We conclude that Ser352 is more dominant in promoting condensation than Ser341. Combining phosphorylation of Ser356 with phosphorylation of Ser341 (**2Pa**) or with phosphorylation of Ser352 (**2Pc**) resulted in no condensates, like the effect of Ser356 phosphorylation alone (**1Pc**) (Figure 5C). We conclude that Ser356 phosphorylation prevents condensate formation even in the presence of the oligomerization–activating phosphorylation at Ser341 or Ser352. However, when the 3 phosphorylations were combined (**3P**), we observed condensate formation despite the phosphorylation at position 356. This shows that a combination of both Ser341 and Ser352 phosphorylations overcame the inhibitory effect of phosphorylating Ser 356. Using a peptide library with all possible combinations of phosphorylations makes it possible to elucidate both the role of each phosphorylation in controlling the oligomerization processes of Tau. Moreover, it enables demonstrating the synergism between the different phosphorylation sites. The library thus provides a detailed mechanistic insight about how the combinations of the phosphorylations contribute to the oligomerization processes.

Moreover, our results revealed that phosphorylation of Ser352 is highly important for Tau aggregation. Phosphorylated Ser352 can be found in a type of Tau filament, named PHF, which was isolated from brains of AD patients^47^. However, it could not be phosphorylated *in vitro* by kinases combination including PKA, GSK, PKA+GSK, P12, P20, PKA+GSK3b and PKA+SAPK4^13, 47-49^. This suggests that phosphorylation of this site *in vitro* is challenging, explaining why this site was not studied extensively in previous research. Working at the peptide level, using our advanced phosphorylated peptide synthesis methods, allowed us to pinpoint Ser352 as crucial site for Tau aggregation.

### The effect of the phosphorylated peptides on the aggregation of Tau-RD

Studying biological systems as closely as possible to their native forms is ideal but creating such a relevant library of phosphoproteins is currently not feasible. Studies at the peptide level enabled us to determine the exact contribution of the individual phosphorylation sites and the combinations between them to Tau oligomerization. We opted on the combination between synthetic phosphopeptides library and recombinant Tau-RD to study the effect of phosphorylation on aggregation at the protein level. This domain was selected because it is the one that drives the amyloid formation in brains of AD and CTE patients.

### The micro-environment of the phosphorylation sites is responsible for controlling Tau R4 oligomerization

SASA calculations of the 29 models of Tau R4 show that pSer352 is significantly less exposed to the solvent compared with pSer341 and pSer356. In the calculated conformations, pSer352 is buried and is forming hydrogen bonds with the backbone of Val337 (e.g Figure 4D model 2). A closer examination of model 1 shows that pSer341 and pS356 are located at the peripheral parts of the peptide with the closest interactions being at distances larger than 4 Angstroms (Figure 4D). This suggests that these phosphate groups do not bind any backbone or side chain residues within the peptide (Figure 4E). Ser341 and Ser356 still differ in their micro-environments. Ser341 is in the IDR of Tau 336-358, while Ser356 is located within the IDR tail, adjacent to the structured domain of the peptide. The residues surrounding Ser341 are the charged Lys340 and Glu342, while the residues surrounding Ser356 are the small Gly355 and the hydrophobic Leu357.

The microenvironments described above for each phosphorylation site explain the observed effects of the different phosphorylations on the oligomerization of the Tau R4 peptide. Phosphorylation of Ser352 resulted in phosphoserine which is less accessible to water and salt. Thus, the peptides including this phosphorylation, **1Pa**, **1Pb**, **2Pa** and **3P,** are less soluble and more available to form interactions with additional peptides or proteins (such as Tau-RD) resulting in condensation or aggregation. Phosphorylation of Ser356, as in **1Pc**, did not show any type of oligomerization. This can be explained by a possible interaction of the phosphate group with water molecules, which leads to better solubility of the entire peptide. Our suggested model correlates with the experimental findings regarding the activating and inhibiting phosphorylation.

High salt concentration inhibited condensate formation by **1Pa**, containing phosphorylation only at Ser341, but not by peptides containing phosphorylation at position 352. This correlates well with our model and can be explained by the higher exposure of Ser 341 to the environment and thus to the salt, which at high concentrations can inhibit the molecular interaction leading to condensation. The micro-environment of the peptides can also explain why **1Pa** formed smaller condensates than **1Pb**. The same principles also explain why 1Pa condensates are smaller and the peptide is concentrated at the boundaries of the condensate. The buried pS352 was not reactive enough to initiate aggregation on its own (it initiated condensation) but it did induce Tau-RD aggregation. pS356 on the other hand is too exposed to allow participation of the peptide in aggregation events with or without Tau-RD.

### Phosphorylated peptide libraries as tools to study defined patterns in proteins

Analyzing the effect of phosphorylation pattern dependent processes in proteins requires using numerous complementary biophysical and biochemical *in vitro* analyses. This, in turn, requires large quantities of high-purity labeled and non-labeled multiphoshporylated peptides since working at the protein level is impractical. As the standard SPPS strategies fail to produce such peptides in sufficient quantities and speed, the synthesis part remains the limiting factor in elucidating phosphorylation dependent processes such as oligomerization. Our results show that applying our newly developed accelerated SPPS method to rapidly synthesize a library of a well-characterized Tau derived multiphosphorylated peptides enabled such in depth analysis of the R4 domain oligomerization.

In summary, our results reveal key insights about Tau aggregation that could not be obtained at the protein level, given the limitations of the existing methods. These insights include the critical role of Ser352 phosphorylation in promoting Tau aggregation and the inhibitory effect that Ser356 has on Tau aggregation. These results provide starting points for further studies at the protein level in the future, when technology will enable such studies. Our studies provide a methodology for systematically studying the effect of phosphorylation patterns on proteins. We suggest to overcome the problems that limit the studies at the protein level by a hybrid peptide-protein approach. We demonstrated that the synthetic phosphorylated peptide libraries, which were obtained in high quantities of pure compounds, enable the use of diverse biophysical methods for gaining insights on Tau oligomerization processes. Such approaches can be applied in the future for other proteins and other PTMs, bridging the gap in the protein level studies of the specific effects of PTM patterns.

## Methods

### Peptide synthesis, labeling and purification

The peptides were synthesized on a rink amid resin with a 0.48 mmol/gr loading after 30 min of swelling in DMF. All peptides were synthesized using the Stirring-based accelerated peptide synthesis approach ^20^. A reactor containing a sintered glass filter, and a heating jacket connected to a circulating 90 °C water bath. The reactor was equipped with an overhead 5-fin turbine PTFE impeller. Solvents and reagents were inserted directly by a feeding line and drained by vacuum filtration. Each cycle started with Fmoc deprotection followed by a 5 mL DMF wash, continued with 1 min coupling and a second wash. Fmoc-Ser/(HPO3Bzl)-OH and its sequential residue were double coupled. 3 mL of coupling mixture containing 3 equiv of protected amino acid, 2.9 equiv of HATU, and 8 equiv of DIEA was added. Fmoc deprotection was done by adding 3 mL of a 0.5% DBU solution in DMF for 10 s. Peptides were cleaved using a freshly prepared TFA cocktail, and were purified by Waters preparative HPLC, using a reverse-phase C18 column with a TDW/ ACN gradient. All peptides were characterized by electron spray ionization mass spectrometry (ESI-MS) (Figure S1). Peptide purity was tested by dissolving 80 µL of peptide in water injecting into analytical HPLC. Peptides were fluorescein labeled For FRAP and fluorescence microscopy experiments, with 5(6)-carboxy fluorescein at their N’ termini. Fractions containing the desired phosphorylated peptides were lyophilized and the peptides were used without further purifications.

### Turbidity assay

150 μL **0P** and **3P** (with 5% FL-peptide each) were tested at peptide concentrations of 50, 300 and 1000 µM. The solutions contained 25 mM phosphate buffer at pH of 7.4 and 10 % PEG 400 (buffer A) with NaCl concentrations of 0, 150, 300 and 1000 mM. Triplicates were measured at λ=350 in BioTek Synergy H1 plate reader.

### Fluorescence microscopy

150 μL **0P** and **3P** (with 5% FL-peptide each) at peptide concentrations of 50, 300 and 1000 µM were also studied using fluorescence microscopy. 10 µL of 50 µM peptide from each sample were applied onto a glass slip and visualized under Zeiss Axio scope A1 microscope with a green-fluorescent filter. Images were taken with an AxioCam ICc 3 camera, using Zen software.

### Fluorescence recovery after photobleaching (FRAP) experiments

FRAP experiments were performed using an FV-1200 confocal microscope (Olympus, Japan) equipped with a sim scanner which enables bleaching while scanning. All experiments were done using a 60x/1.42 oil objective. Imaging was done using the 488 nm laser line and emission was collected 500-5400 nm. 5 % of **FL-3P** and 95 % of **3P** were dissolved in 100 µL buffer A to final concentration of 25 mM phosphate buffer and 150 mM NaCl and 50 µM peptide. The measurement started by adding 10% (V/V) PEG-400 and samples were bleached after 15 and 45 min to evaluate the maturation of condensates. A specific region in the sample was exposed to 100 % laser intensity for 120 msec. The interval between image scans was 2 sec. Photobleaching correction and recovery were calculated using the OriginPro 2022 software. Final FRAP recovery curves are the average from measuring three different condensates.

### CD of 0P and 3P

120 µM of 0P and 3P, in a 25 mM phosphate buffer at pH of 7.4 and 150 mM NaCl CD spectrum were determined using JASCO J-810 spectropolarimeter and Spectra manager software. The temperature was 25 C° and controlled by Peltier thermostat, in a 0.1 cm quartz cuvette for far-UV CD spectrometry, ranged from 190 to 260 nm.

### Aggregation assay

To examine the aggregation properties of the peptides, 100 µM of peptides were incubated with and without 10 µM Tau-RD and 20 mM phosphate buffer at pH of 7.4 and 150 mM NaCl. 150 µL of the final solutions were incubated in a temperature of 37 C° for 10 days. The solution was then dried on silicon wafers and examined using SEM Apero 2S LoVac. Each experiment was repeated between three to nine times.

## Conflict of Interest

The authors declare no conflict of interest

## Acknowledgments

MH is supported by the European Innovation Council (EIC) under the European Union’s Horizon Europe research and innovation program (No. 101046369) and by the Israel Science Foundation grant no. 1805/22. SGDR was supported by grants of the Campaign Team Huntington and Alzheimer Nederland (No. WE.03-2019-03). He is PIs of the gravitation consortium FLOW, funded by the Nederlands Minister of Education, Culture and Science.. AF thanks the Minerva Center for Bio-Hybrid complex systems and the Saerree K. and Louis P. Fiedler Chair in Chemistry. This study was supported by a visiting professorship in HUJI (to SR). We thank Dr Ofrah Faust and Prof Rina Rosenzweig from the Department of Chemical and Structural Biology, Weizmann Institute Israel for generously contributing the Tau-RD protein.

## Notes

### Competing Interest Statement

The authors have declared no competing interest.

### Summary of Updates

The resolution of the figures was improved in the revised vesrion

