## Supplementary figures for "The role of specific phosphorylation patterns in the oligomerization of Tau-R4"

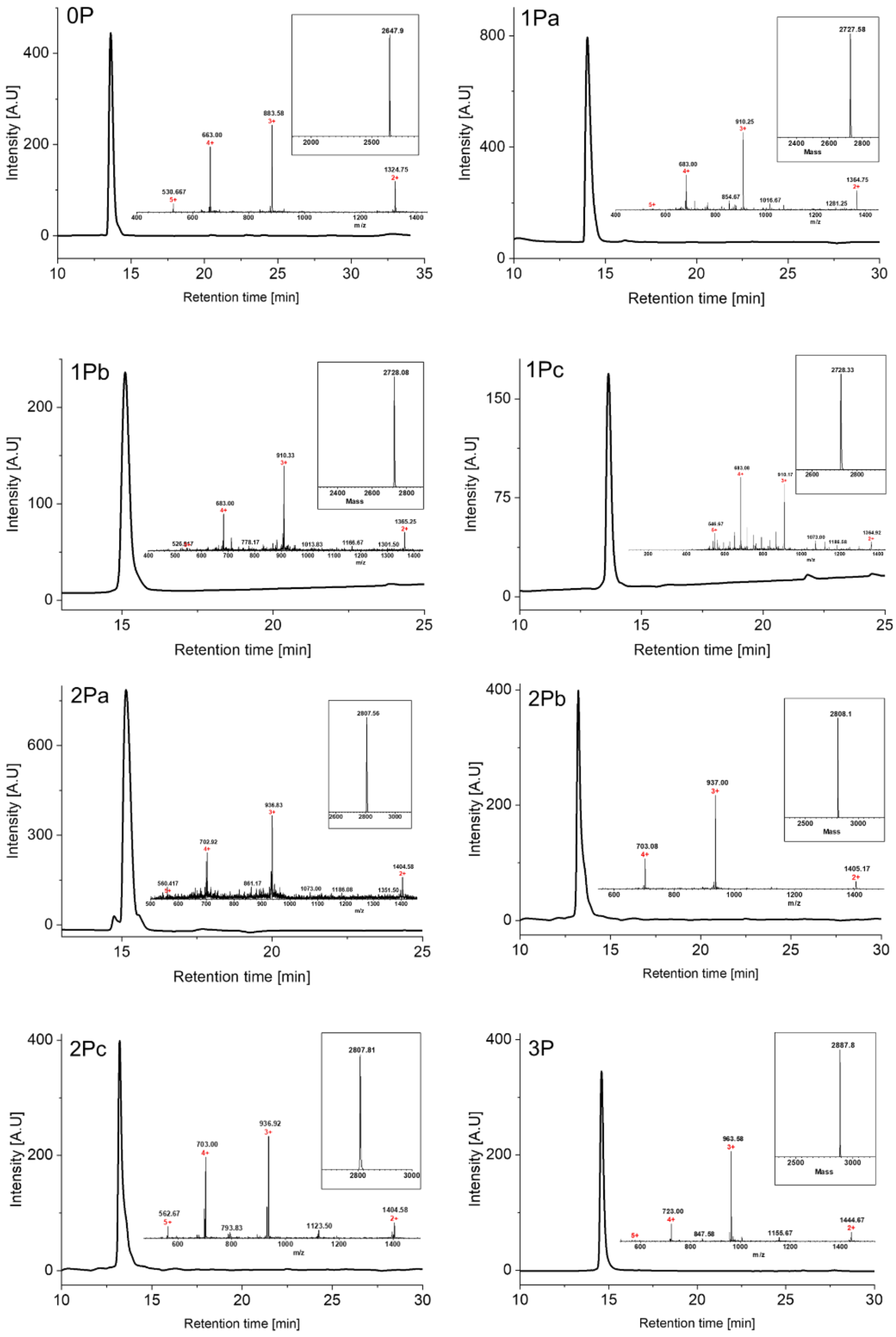


**Fig. S1.** Analytical HPLC chromatograms and mass spectra. (insert in each panel) of the peptides synthesized in the current study. The calculated mass of **0P** is 2648.0 Da, the observed mass is 2647.9 Da. The theoretical mass of the mono-phosphorylated peptides are 2728.0 Da. the observed masses are: **1Pa** = 2727.6 Da, **1Pb** = 2728.1 Da and **1Pc** = 2728.3 Da. The calculated masses of the di-phosphorylated peptides are 2808.0 Da. The observed masses are: **2Pa** = 2807.6, **2Pb** = 2808.1 and **2Pc** = 2807.8. the theoretical mass of **3P** is 2887.9 and the observed mass is 2887.8. overall, HPLC chromatograms show that all peptides are >95% pure.

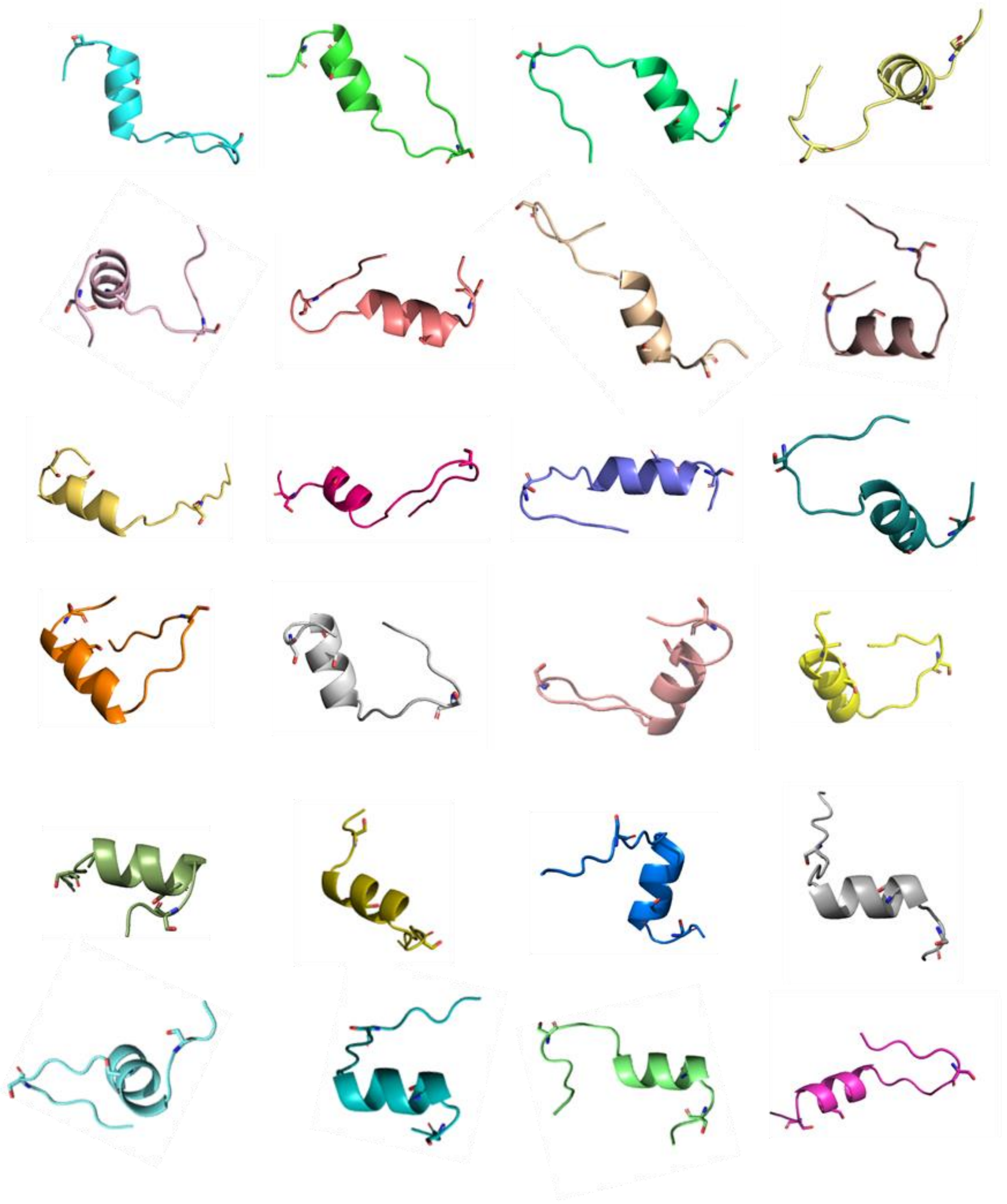

**Fig. S2.** The 24 models of Tau 336-358 that were generated by PEP-FOLD3 [41] and used for molecular surface and SASA calculations. The 3 serines are represented by sticks. In all models Ser 341 is located in the disordered domain, Ser352 in the helical structured domain and Ser356 in the IDR rail,
